# Adaptive cognitive maps for curved surfaces in the 3D world

**DOI:** 10.1101/2021.08.30.458179

**Authors:** Misun Kim, Christian F. Doeller

## Abstract

Terrains in a 3D world can be undulating. Yet, most prior research has exclusively investigated spatial representations on a flat surface, leaving a 2D cognitive map as the dominant model in the field. Here, we investigated whether humans represent a curved surface by building a dimension-reduced flattened 2D map or a full 3D map. Participants learned the location of objects positioned on a flat and curved surface in a virtual environment by driving on the concave side of the surface (Experiment 1), driving and looking vertically (Experiment 2), or flying (Experiment 3). Subsequently, they were asked to retrieve either the path distance or the 3D Euclidean distance between the objects. Path distance estimation was good overall, but we found a significant underestimation bias for the path distance on the curve, suggesting an influence of potential 3D shortcuts, even though participants were only driving on the surface. Euclidean distance estimation was better when participants were exposed more to the global 3D structure of the environment by looking and flying. These results suggest that the representation of the 2D manifold, embedded in a 3D world, is neither purely 2D nor 3D. Rather, it is flexible and dependent on the behavioral experience and demand.

## 1. Introduction

People often rely on printed or electronic maps for everyday navigation. Due to the 2D nature of these formats, most maps are simplified 2D representations of the 3D world in which we live. The maps often lack information about the elevation of undulating terrain (except for the contour plot) and give us the impression that both the physical map as well as our internal representation of the world is mainly 2D. The vast majority of research on spatial navigation and cognitive mapping has also been conducted on horizontal, flat, 2D surfaces, despite real environments being more complex (Boccia et al., 2014; Moser et al., 2017). Whether it is sufficient to have a 2D surface-based map or whether humans build a full 3D model of the world is an important question that is not fully understood.

Some of the early behavioral and neurophysiological studies suggest that surface-dwelling animals such as rodents and humans have a 2D or at best 2.5D representation of the environment, as their movements are constrained to the earth’s surface due to gravity (Jeffery et al., 2013). Humans and dogs showed better within-floor spatial memory than across-floor memory in a multi-level building (Brandt & Dieterich, 2013; Hölscher et al., 2006; Montello & Pick, 1993). In the brain, head direction cells serve as a compass system and these cells are mainly sensitive to the horizontal component of the heading (azimuth) but not to the vertical pitch (Stackman & Taube, 1998). Head direction cells encoded the direction, relative to the local plane of locomotion, even when the locomotion plane was rotated in 3D space such as with a vertical wall or ceiling (Calton, 2005; Page et al., 2018; Taube et al., 2004, 2013). Rather than showing a 3D volumetric receptive field, entorhinal grid cells have shown similar firing patterns on a horizontal plane and its connected slope, as if the slope was an extension of the horizontal surface (Hayman et al., 2015). Based on neuroscientific findings, Jeffery et al. proposed that 3D space is encoded as a mosaic of local planar maps (Jeffery et al., 2013).

On the other hand, it is conceivable that even surface-residing animals could build a more complete 3D model of the environment than a mere surface model. Those who navigate on undulating terrain must consider the elevation to minimize the energy cost of moving upwards. This has been shown in a study where people tried to avoid local hills, taking a detour when they were asked to take the shortest route while solving the travelling salesman problem in a non-flat environment (Layton et al., 2010). Furthermore, a slope can be utilized as a salient orientation cue for spatial memory tasks in humans (Holmes et al., 2015; Nardi et al., 2011, 2021; Steck et al., 2003), rats (Grobéty & Schenk, 1992; Wilson et al., 2015), and pigeons (Nardi & Bingman, 2009). It is also reported that the hippocampus in both rodents and humans encodes horizontal as well as the vertical spatial information similarly well when subjects move around in semi-volumetric environment (Grieves et al., 2020; Kim et al., 2017). Similarly, grid cells with 3D ellipsoidal receptive fields were observed in rat and bat entorhinal cortex when these animals explored an open volumetric space (Ginosar et al., 2021; Grieves et al., 2021).

A simplified surface-based map enables efficient and compressed encoding of the world, whereas a fully volumetric 3D map would allow flexible route planning beyond the current plane of locomotion. It remains unknown which type of the representation is utilized by humans who navigate on a curved surface embedded within a 3D world. To tackle this question, we built a novel 3D virtual environment composed of a flat and curved surface and asked participants, with varying degrees of movement, to learn the location of objects on the surface in the series of online behavior experiments (Experiment 1: driving alone, Experiment 2: driving with additional vertical viewing, Experiment 3: flying). We then asked participants to retrieve either path distances on the surface or Euclidean distances in 3D from their memory. Distance estimates for the flat and curved parts were analyzed to probe which type of cognitive maps participants built.

## 2. Experiment 1

### 2.1. Introduction

In this experiment, participants learned the location of objects on seamlessly connected flat and cylindrical surfaces by driving with their heads parallel to the tangent of the surface (Fig. 1). Crucially, the objects were positioned in a way that between-object path distances were identical on the flat and curved surfaces (e.g. AB=AG), whereas the Euclidean distance was shorter for the curved surface (e.g. AB<AG). Given that all objects and participant views and movements were restricted to the local surface, it could be sufficient to remember the object location relative to the boundary of the surface, treating the curved surface as an extension of the flat surface and disregarding the position of the surfaces within the global 3D world. Thus, one might develop a flattened 2D map as illustrated in the schematic figure Fig. 1B. In this case, there should be no systematic bias in path distance estimation for the objects on the flat and curved areas (e.g. AB=AG). Further, participants should show difficulty in estimating 3D Euclidean distance if they are asked to do so later. On the other hand, participants might also automatically encode the 3D global layout of the environment and build a volumetric 3D map (Fig. 1A), even if it was not strictly necessary for the object location learning task. In this case, participants would be good at the Euclidean distance estimation task and 3D map knowledge might even interfere when participants try to recall the path distance, resulting in an underestimation bias for the curved path (e.g. AB<AG). We tested these predictions in an online virtual reality (VR) experiment.

**Figure 1.**
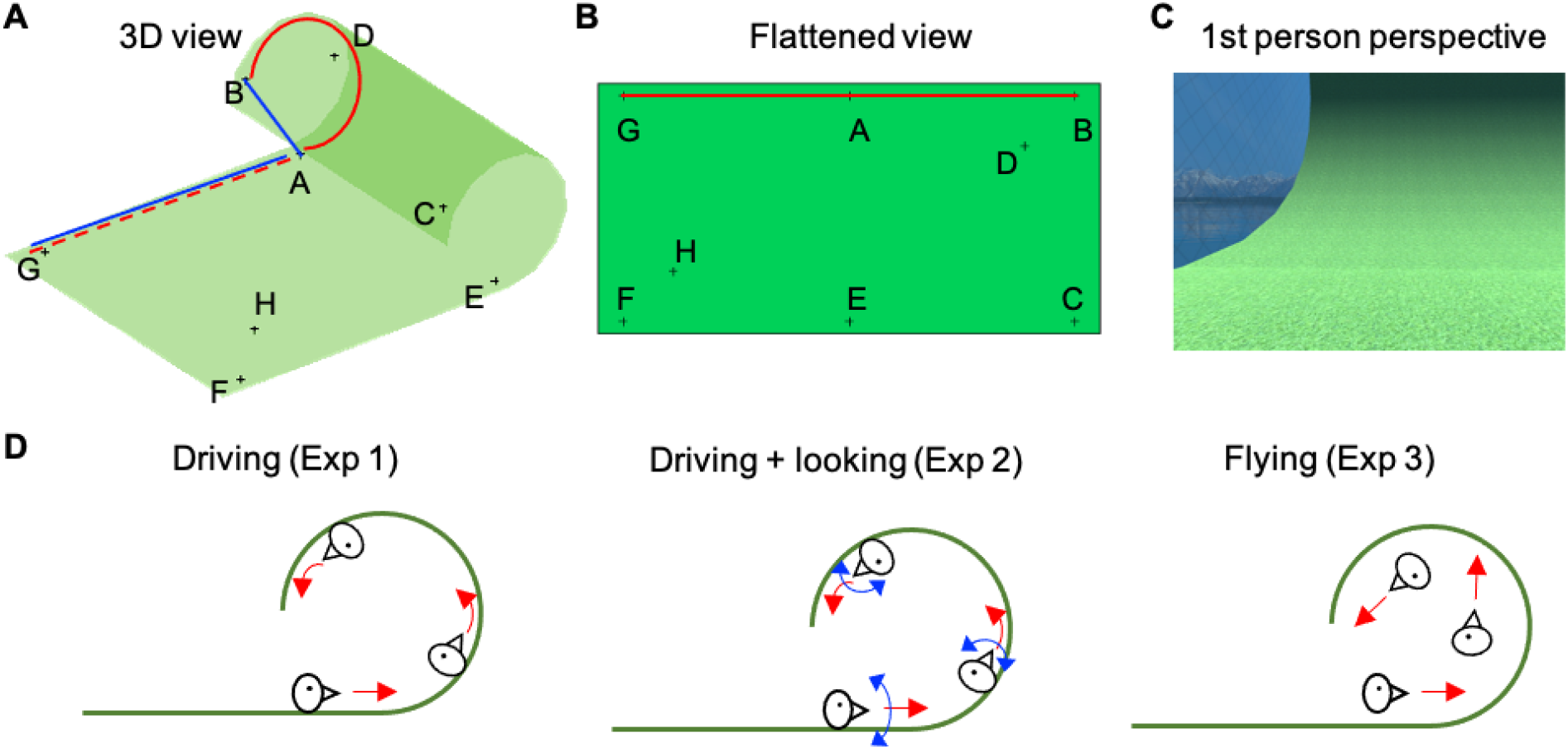
The virtual environment and exploration methods used in the three experiments. **A**. The virtual environment consisted of a flat square and a curved surface with an arc length of 270 degrees. The length of the square was identical to the arc length of the curve, so that between-object path distances (red lines) were identical on the curved and flat surfaces (e.g. AB=AG). In contrast, 3D Euclidean distances between the objects (blue lines) were shorter on the curve (e.g. AB<AG). The virtual environment also contained semi-transparent walls at the rim of the surface which prevented participants from moving beyond the surface (not shown here for visibility). **B**. The same environment can be represented in an abstract, flattened 2D map. **C**. Participants explored this environment from a first-person-perspective. **D**. Schematic figures illustrating the movement method. In Experiment 1 participants drove on the surface and always remained parallel to the tangent of the surface. In Experiment 2 participants could also look up and down while driving. In Experiment 3, participants could freely fly.

### 2.2. Method

#### 2.2.1 Participants

Participants were recruited from the online experiment platform (www.prolific.co). Forty-two participants (female = 12, mean age = 22.7 ±4.0 years) were in the path distance group and 45 participants (female = 16, mean age = 24.2 ±4.8 years) were in the Euclidean distance group. All participants self-reported having no current, or history of, neurological or psychiatric disorder. This study was approved by a local research ethics committee.

#### 2.2.2 Virtual environment and the movement within it

The virtual environment was composed of a flat square and a curved surface (Fig. 1A). The curved section was continuous with the flat section, with an arc of 270°. The entire structure resembled a paper that was being rolled up or a half-pipe in a skate park (Fig. 1A). The long arc was used to maximize the difference between the 2D path distance and the 3D Euclidean distance between the two extreme ends of the arc. The arc length (25 virtual units) was identical to the length of the flat surface. A green grass texture was applied to the whole surface and semi-transparent walls formed the boundary of the surface, preventing participants from moving beyond the surface. A snowy mountain and lake landscape was used as a background. We used Unity 2018.1.9f2 (Unity Technologies, CA) to implement the virtual environment and deploy the experiment into the web browser.

Participants explored the virtual environment from a first-person perspective using a desktop-based VR. Participants were told that they were driving a vehicle on the surface and that the vehicle could not fall off. They could move forward/backward and turn the vehicle right/left using the arrow keys. Participants’ heading (i.e. viewing) direction was always parallel to the tangent of the surface. Their view was rather limited on the curved portion of the surface due to the concavity of the environment. The speed of the movement was constant throughout the environment (8 virtual units/sec). More specifically, there was no acceleration or deceleration due to gravity and participants could move similarly on the curved and flat parts of the environment. Snapshots of the environment from a participant’s perspective are shown in Fig. 1C and Fig. 2A. An example trajectory is shown in <u>Supplementary Video 1</u>. Readers can also explore the virtual environment in the interactive webpage (http://misunkim.github.io/curveSpace_demo).

**Figure 2.**
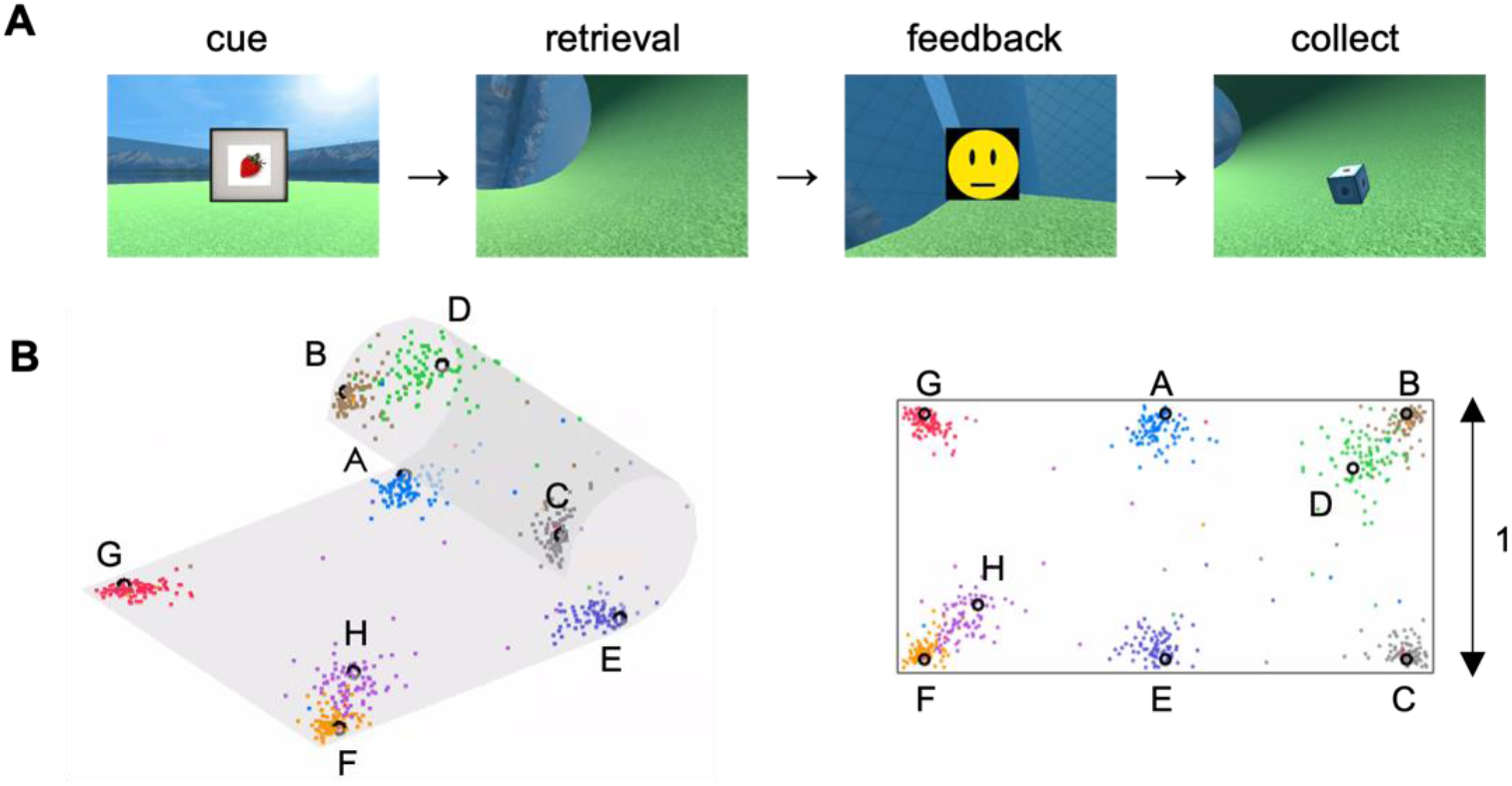
The object-location memory task. **A**. In each trial, a picture cue was shown at the beginning and participants moved to the remembered location of the object (retrieval). After the feedback, participants collected the object, which reappeared at the correct location. **B**. Most participants remembered the location well at the end of the test phase (mean distance error < 0.1) and the distance errors for the objects on the curve (B, C, D) and the flat (F, G, H) sections were not different. Colored dots indicate the remembered location of objects in the last trial for all participants; black circle indicate the true object location. Left, 3D view; right, flattened view. Distance was normalized to the length of the short axis of the surface.

#### 2.2.3 Tasks and analysis

Participants completed the tasks in the following order: familiarization, object-location learning and testing, distance estimation tasks, and debriefing. The whole experiment took about 30 minutes.

##### 2.2.3.1 Familiarization

Participants first familiarized themselves in the virtual environment and practiced the movements. They were instructed to move to a traffic cone. Once they arrived at the cone, another cone would appear, at a new location, and participants had to find the new cone. To help participants to quickly find the cone, which could be invisible due to curvature of the environment, a guide arrow was shown on the ground throughout the whole familiarization period.

##### 2.2.3.2 Object location learning and test phase

Following the familiarization period, participants learned the locations of 8 objects in the environment (Fig. 2). The objects were cubes with pictures of animals or fruit on their faces. The 8 locations were constant, while the assignment of each picture cube to each location was randomized across participants. Each picture cube appeared once sequentially and participants were guided to move to each picture cube.

At the beginning of each trial in the test phase (following the learning phase), a picture cue was presented at the center of the screen and the participant was teleported to a random starting location. They then moved to the remembered location of the cued picture cube and pressed the spacebar. There was a time limit of 60 seconds per trial. Very few trials (0.1 trial per participant on average) were aborted due to the timeout. After each trial, participants received feedback on their displacement error (5 scales from a frowning to a smiling face) and were shown the correct location of the picture cube. They had to move to the correct location for the next trial to begin. Each object was tested between 4 and 8 times until the participant reached the learning criteria (distance error of less than 25% of the short axis of the environment). For the distance estimation analysis, we only included the participants who remembered all 8 objects well (i.e. distance error in either of the last two trials was less than 25%).

##### 2.2.3.3 Distance estimation tasks

We then asked participants to estimate the between-object distances outside the virtual environment. The path distance group was instructed to estimate the distance by imagining how long it would take to move from one picture cube to the other picture cube (e.g. red lines on the surface in the schematic Fig. 1A). In contrast, the Euclidean distance group was instructed to imagine a straight line between the object in 3D space (e.g. blue lines in Fig. 1A). To ensure that the participants had understood the definition of a 3D Euclidean line, we showed a short video that visualized it from a first-person-perspective in the virtual environment (<u>Supplementary Video 2</u>). This led to the possibility that participants would gain additional knowledge of the 3D environment from this short video instruction, specifically in the Euclidean group, even though their movement and views were restricted to the surface beforehand. In order to minimize the additional exposure to the 3D structure via the instruction video, we made the video short (less than a minute) and didn’t show the picture cube during this instruction, thus preventing participants from further learning the distance between the objects. Importantly, a bird’s eye view of the 3D environment like Fig. 1A was not shown to participants and we did not inform them about the upcoming distance estimation tasks while they were learning the object locations inside the virtual environment. Therefore, participants must have reconstructed a cognitive map of the virtual environment from their memory to estimate either path or Euclidean distance upon request.

The knowledge of distance was probed with two types of tasks: a comparison task and a slider task. In the comparison task, a triplet of objects was presented on the screen and participants had to decide whether the object on the top row was closer to the 1^st^ or 2^nd^ object on the bottom row (Fig.3A). Participants did not receive performance feedback. The key triplets were the comparison between the path on the flat and curved surface (AB vs. AG, or EC vs. EF, see the location naming in Fig. 1A) with each being presented four times (8 trials in total). The true path distance was identical for the curve and linear sections and therefore neither path should be reported as shorter (i.e., closer) above chance level (50%) if participants had an accurate surface map and were able to retrieve distances on that map. On the other hand, if the knowledge of the 3D nature of the map and Euclidean distance interferes with the surface map knowledge, participants might show an underestimation bias for the curve path, because the Euclidean distance was shorter for the objects on the curve. As a manipulation check, we also included trivial triplets that could easily be solved by either path or Euclidean distance (e.g. FH vs. FG, BD vs. BA) and the triplets where both path and Euclidean distances were identical. After excluding a few outliers (see Results section), whose accuracy was chance level for the abovementioned easy trials, we tested whether the response rate for choosing the curve distance as shorter than the linear distance was above chance using a Wilcoxon-signed rank test due to non-normality of the data.

**Figure 3.**
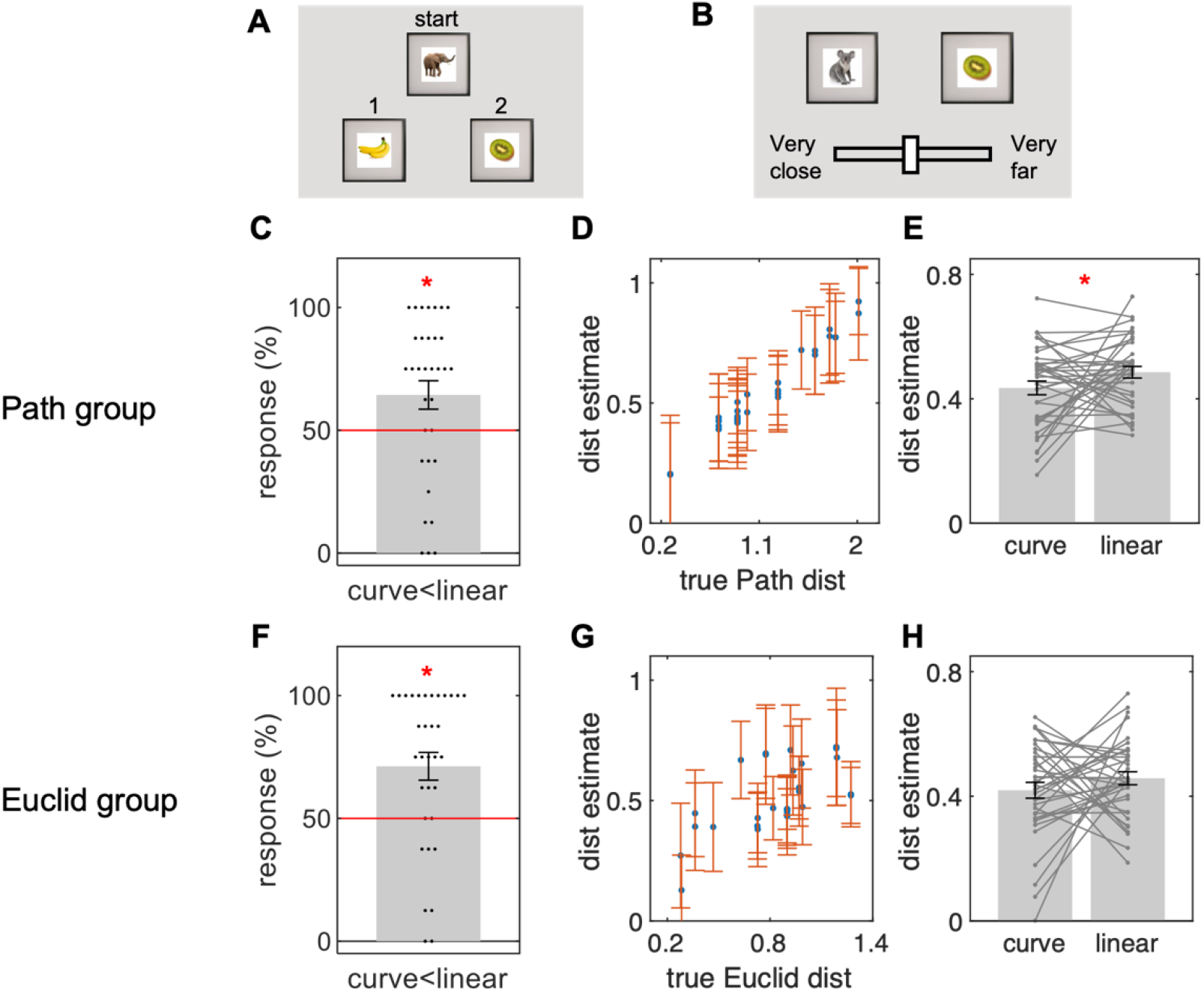
Distance estimation task in Experiment 1 (driving). **A**. In the comparison task, participants chose whether the first or second object was closer to the top object from object triplets. **B**. In the slider task, participants reported the relative distance of all object pairs using the continuous scale.**C-E**. Results from the path group. **C**. There was an underestimation bias for the curve path, as shown by over 50% of responses choosing the curve path as shorter than the linear path. Black dots = individual participants; red line = chance level. **D**. In the slider task, participants’ estimated path distances were highly correlated with the true path distances. Each dot = group mean distance estimate for the unique object pair. **E**. The mean path estimate for the curve was slightly but significantly shorter than the linear one. Dots = individual participant’s distance estimate between 0 (“very close”) and 1 (“very far”). **F-G**. Results from the Euclidean group. **F**. Participants correctly reported the curve distance as shorter than the linear distance during the comparison task. **G**. In the slider task the estimated distances were positively correlated with true Euclidean distance. **H**. Curved distance estimate was not significantly different from the linear distance. All error bars are group SE.

In the slider task, participants rated the distance between pairs of objects by adjusting the slider bar with a range from “very close” to “very far” (Fig. 3B). We scaled the slider bar between 0 (“very close”) and 1 (“very far”). All possible pairs of objects were presented twice (8 objects, 28 unique pairs, 56 trials in total) and we averaged the two ratings for each unique pair of objects within participant. We first quantified the overall consistency between the subjective distance rating and the true path or Euclidean distances using Spearman correlation. We then focused our analysis on the key pairs where the curve and linear distances were maximally dissociated (curve: AB, EC vs linear: AG, EF). We tested whether the curve distance was rated shorter than the linear distance using a paired t-test.

##### 2.2.3.4 Debriefing

After the main tasks, participants completed the short debriefing forms. The questionnaire included the question on whether they had paid attention to the distance or size of the virtual environment during the object-location tests and what types of strategies they used during the task (e.g. first-person perspective or 3D bird’s eye view or flattened map). We report the percentage of self-reported strategies for the distance estimation task from the participants who met the object-location learning criteria.

##### 2.2.3.5 Statistical analyses

All analyses were conducted in MATLAB and we report the group mean ± standard deviation, degrees of freedom, and p-values, unless stated otherwise.

#### 2.2.4 Control experiments

When participants drive on the surface, the view point is above the surface with certain eye height. The path length at the eye height is shorter than the path length at the surface level in the concave surface like the one used in the current experiment. We asked participants to estimate the path length between the picture cubes which were lying on the surface, therefore participants should estimate the path length at the ground level, not at the eye height level. Nevertheless, to rule out the possibility that participants underestimated the path length on the curve section due to mistaking the definition of path length, we conducted a control experiment which manipulates the eye height. We increased the eye height from 1 virtual unit to 2.5 virtual unit in this control experiment (Supplementary Fig. 1). If participants measured the distance between the objects at the eye level, we would expect a significantly stronger underestimation bias for this new experiment. Similar to the main experiment, a new group of 44 participants were recruited and estimated the path distance (female = 21, mean age = 22.8 ±3.7 years). On a side note, we also increased the size of the traffic cone and picture cubes so that these objects were easily visible at the new eye height.

In the second control experiment, we built a convex environment where participants drive on the outside of the cylindrical surface (Supplementary Fig. 2, <u>Supplementary Video 3</u>). In the new environment, path length at the eye level is longer on the curved section. Importantly, the emotional factor and the accessibility of Euclidean shortcut also differ from the main experiment. People can feel a stronger sensation of being upside down and falling when they drive outside of the curve compared to when they drive inside of the curve. More crucially, Euclidean shortcut might become less imaginable in the convex setting because that would require a building of tunnel through the surface, in comparison to relatively easily imaginable bridge or flying route in the concave setting. We tested whether there is still an underestimation bias for the path along the curved section in the convex setting with a new group of 45 participants (female = 22, mean age = 23.0 ±3.8 years).

### 2.3. Results

#### 2.3.1 Object location memory result

Most participants remembered the location of all 8 objects well (mean repetitions per object = 5.2 ±0.9, mean distance error on the last two trials = 0.09 ±0.09 for all 87 participants. All distances are reported as relative size to the short axis of the environment, Fig. 2B). We had to exclude 13 participants from further analysis who did not meet our learning criteria (see Method 2.2.3.2). The final distance errors for the objects on the curved part of the environment were not significantly different from those on the flat part (mean error for the objects B, C, and D on the curve = 0.06 ±0.02 vs. object F, G, and H on the flat = 0.06 ±0.02, t(73) = 0.7, p = 0.49). This suggests that participants had comparably good memory for the flat and curved surfaces.

Of note, we found a small bias in participants’ memory for the objects at the intersection between the flat and curved sections (location A, E). The remembered locations for these middle objects were slightly skewed to the flat side of the environment, such that they were placed slightly closer to the object at the end of the flat part rather than the object at the end of the curved part. This was due to the tendency of participants to approach the middle object location from the flat side rather than the curved part, where the view was inherently limited due to the curvature. The movement trajectories are visualized in Supplementary Fig. 3. The resulting difference in the between-object distance for the linear and curved parts was very small, but significant (curve = 0.93 ±0.06 vs. linear = 0.87 ±0.07, t(73) = 5.4, p < 0.001).

#### 2.3.2 Distance estimation result for path group

Following the object-location test, the path group participants were asked to estimate the between-object path distances on the surface. Key pairs of objects had been placed at equal distances on the curved and flat sections, and we tested whether there was a bias in the distance estimation that could be explained by interference from 3D map knowledge.

During the comparison task, most participants (33 out of 37) passed the manipulation check (e.g. above chance level accuracy for very easy trials, see Method 2.2.3.3). Critically, in the key comparison trials, where participants had to choose between the equidistant options of the curved and linear paths, they showed a significant underestimation bias for the curved path. This is indicative of an interference effect from the 3D map knowledge (mean rate = 64 ±33%, Wilcoxon sign-rank test, n = 33, Z = 2.2, p = 0.013, 1-tailed, Fig. 3C).

We then examined the distance ratings from the slider bar task where participants indicated the perceived distances of all object pairs using the continuous scale ranging from “very close” (0) to “very far” (1). Participants’ path distance estimates were highly correlated with the true 2D distances (ρ = 0.69 ±0.26, Fig. 3D), implying that most participants had an overall good path distance estimation ability. Importantly, the curve distance estimate was again slightly but significantly shorter than the linear distance (mean estimate for curve = 0.44 ±0.13, linear = 0.49 ±0.11, paired t-test, t(36) = 1.7, p = 0.045, one-tailed, Fig. 3E). Thus, in both the slider bar and comparison tasks we found consistent evidence for an influence of the 3D curve on distance estimates.

At the end of the experiment participants were asked about their strategy during distance estimation. The option “I imagined myself moving from one picture to the other picture and estimating the distance (first-person perspective)” was selected by 24% (Fig. 1C), 57% selected “I had the map of the 3D curved environment in my mind and measured the distance on that map (bird’s-eye view)” (Fig.1A), and only 19% selected “I had the map of the environment in my mind, but it was rather a simple, flat map.” (Fig.1B).

In the control experiment where the eye height was increased, we found a similar magnitude of the underestimation bias for the curved section (Supplementary Fig. 1), implying that participants did correctly understand the definition of path length at the surface, rather than mistakenly reporting the path length at the eye height. In other control experiment where the convex surface was used, we no longer saw the underestimation bias for the curved section (Supplementary Fig. 2). Similar to the main concave experiment, a small number of participants reported to use a flattened map strategy (c.f. first-person perspective = 38%, 3D map = 35%, flattened map = 27%).

#### 2.3.3 Distance estimation result for Euclidean group

The Euclidean group were asked to recall 3D Euclidean distances instead of path distances on the surface. In the comparison task, 33 out of 37 participants solved the easy trials with above chance accuracy and these participants correctly chose the curve distance as shorter than the linear distance (71 ±32%, sign-rank test, Z = 3.2, p = 0.001, Fig. 3F). However, the result of the slider task was ambiguous. Participants’ distance estimates were positively correlated with the true Euclidean distance (Fig. 3G, ρ = 0.38 ±0.25). However, the key curve distance estimate was not significantly shorter than the linear distance (mean estimate for curve = 0.42 ±0.15, linear = 0.46 ±0.13, t(36) = 1.1, p = 0.15, one-tailed, Fig. 3H).

### 2.4. Discussion

Overall, participants showed good spatial memory for objects lying on the flat and curved surfaces and they could also later estimate between-object distances from memory well above chance. Crucially, we found that participants estimated the path distances along the curve as slightly shorter even though the true path distances were identical. This distance estimation bias cannot be explained by an imprecise memory for object-location or poor task comprehension because we only included those participants who passed not only the object-location learning criteria but also the manipulation check for the distance task. Moreover, the opposite pattern of result would be expected if a small bias in location memory error was taken into consideration. However, as we described in the object-location memory result section, participants placed the middle object slightly closer to the objects on the flat side than the objects on the curved side. Thus, if this would have influenced the distance estimation, they should have rather underestimated the linear path, not the curved path. The control experiment with eye height manipulation showed that the path at the eye level above the ground could not explain the underestimation bias. Reduced visibility in the curved part also is also unlikely to explain the underestimation bias for the curve path. It is known that people overestimate the distances between objects separated by barriers (Newcombe & Liben, 1982). Our curved environment can be regarded as containing a barrier because participants could not directly view the objects at the other end of the curve, due to the curve itself. Therefore, one would rather expect an overestimation bias for the curved path if visibility matters. Ruling out the abovementioned factors, we interpret the underestimation bias of the curved path as an influence of knowledge of the 3D representation and potential Euclidean shortcut. Participants never saw these 3D shortcuts in the virtual environment, which were thus, behaviorally irrelevant. The self-reports during the debriefing, which showed the dominance of the 3D map over the 2D map, also supports the view that participants acquired 3D knowledge of the environment.

Of note, we didn’t see the underestimation bias for the curved path in the control experiment with a convex surface. Compared to the concave surface where one can relatively easily imagine a Euclidean shortcut between the points on the surface as a form of a bridge or straight-line along the sight, such a shortcut can be more difficult to imagine in the convex setting where a tunnel through the earth should be built and this might be a reason for the difference in path perception between the convex and the concave setting. However, even in the convex setting, only a small number of participants reported to use a flattened 2D map strategy; rather they reported to use a 3D map or first-person-perspective strategy. Therefore, we believe that people have a level of 3D global knowledge in both concave and convex setting.

However, the influence of 3D shortcuts and/or the self-reported strategy for a 3D map does not imply that humans generally build a precise, metric, 3D map knowledge when they drive on a surface. This is rather unlikely because 3D Euclidean distance estimation was far from perfect when participants were explicitly asked to imagine the 3D distance. They correctly selected the curve distance as the shorter one when they were forced to choose between the curve and linear distances during the comparison task, but their distance estimates for the curve and linear distance were not significantly different during the slider task. This suggests that the perceived difference between the two distances was small and that the difference was only detectible when the two were directly compared against each other. The difference was not detectible when participants were asked to estimate each distance separately, across different trials in the slider task. The slider task was inherently more difficult than the direct comparison task. The range of distances that participants had to remember and estimate with the slider was large, from the nearest object pair (e.g. BD) to the farthest-away object pair (e.g. CG), and the key pairs in the middle of the range.

Taken together, the current experiment showed that participants did not build a strictly 2D map that was agnostic to the global 3D structure. There was an influence of the global 3D structure and Euclidean distance. But participants did not form elaborate 3D map knowledge, which also was not necessary for the behavioral demands (i.e. learning the object-location on the surface while driving). This raises the question of whether humans have a fundamental difficulty in constructing 3D maps of the world. We predicted that the way participants learn the spatial layout of the environment (e.g. how many 3D cues are available in the environment or whether their movement is constrained to the surface or not, etc.) would influence the type of map they would build. This hypothesis was tested in the next experiments.

## 3. Experiment 2

### 3.1. Introduction

In Experiment 1, participants were always facing parallel to the surface while moving around. Thereby, the vertical tilt was zero (Fig. 1D). With zero vertical tilt, participants can see everything in front of them when standing on the open flat environment, whereas they can see distal points on the curved surface only when they look up, because the curvature blocks the view. In the current experiment, we allowed participants to freely look up and down while learning the object locations to facilitate the learning of the global 3D structure (Fig. 1D). The object locations and the following distance estimation tasks were identical to the previous experiment. We tested 1) whether there is an underestimation bias for the curve path in the current setup, and 2) whether participants can better estimate the 3D distance.

### 3.2. Method

#### 3.2.1 Participants

Participants were recruited from the online experiment platform (www.prolific.co). Forty-four participants (female = 10, mean age = 23.9 ±4.7 years) were in the Path distance group and 44 participants (female = 13, mean age = 23.7 ±4.8 years) were in the Euclidean distance group. None of them took part in the previous experiment.

#### 3.2.2 Virtual environment

The identical environment was used as in Experiment 1.

#### 3.2.3 Task and analysis

As previously, the experiment was composed of a familiarization phase, object-location learning and test, distance estimation tasks, and debriefing. The only changes were that participants could additionally look up (away from the surface) or down (towards the surface) using the arrow keys, and the guide arrow on the ground, that showed the direction to the target, was removed to encourage active looking behavior. Most participants looked slightly upwards (mean tilt angle = −11 ±9°) when they were on the curved part of the environment and they looked rather straight or slightly downwards when they were on the flat part of the environment (mean tilt angle = 4±7°) during the object location test. An example viewing trajectory and distribution of tilt angle is shown in Supplementary Fig. 4 and <u>Supplementary Video 4</u>. The rest of the analysis was identical to Experiment 1.

### 3.3. Result

#### 3.3.1 Object location memory

Similar to Experiment 1, most participants showed good memory of object locations (mean repetition per object = 5.1 ±1.1, mean distance error = 0.09 ±0.08 for all 88 participants, all distances are reported as relative to the size of the short axis of the environment, Supplementary Fig. 5). Only those who met our object-location memory criteria were included in the distance estimation analysis, leaving us with a final sample of n = 38 for the path group and n = 35 for the Euclidean group. As before, we found that participants placed the center objects slightly towards the flat side of the environment as opposed to the curved side (distance to the flat side = 0.89 ±0.06 vs. distance to the curved side = 0.92 ±0.06, t(72) = 2.6. p = 0.01). Unlike the first experiment, the final distance error for the objects on the curved section was slightly larger than for the flat section (error for the objects on the curve = 0.06 ±0.02 vs. flat = 0.05 ±0.02, t(72) = 2.6, p=0.012).

#### 3.3.2 Distance estimation: path group

As before, a majority of participants showed good comprehension of the distance estimation task: 35 out of 38 participants showed above chance accuracy for easy trials in the comparison task. During the direct comparison between the curved and linear paths, we found a tendency for participants to choose the curve as shorter (62 ±41%, sign-rank test, p = 0.051, Z = 1.6, 1-tailed, Fig.4A), similar to the result of Experiment 1. This suggests an influence of the 3D representation during path retrieval. Overall, participants showed good path distance estimation during the slider task as previously (ρ = 0.65 ±0.34, Fig. 4B). However, the curve path was not significantly underestimated during the slider task (mean estimate for curve = 0.43 ±0.14 vs. linear = 0.45 ±0.10, paired t-test, t(37) = 1.0, p = 0.15, one-tailed, Fig. 4C).

**Figure 4.**
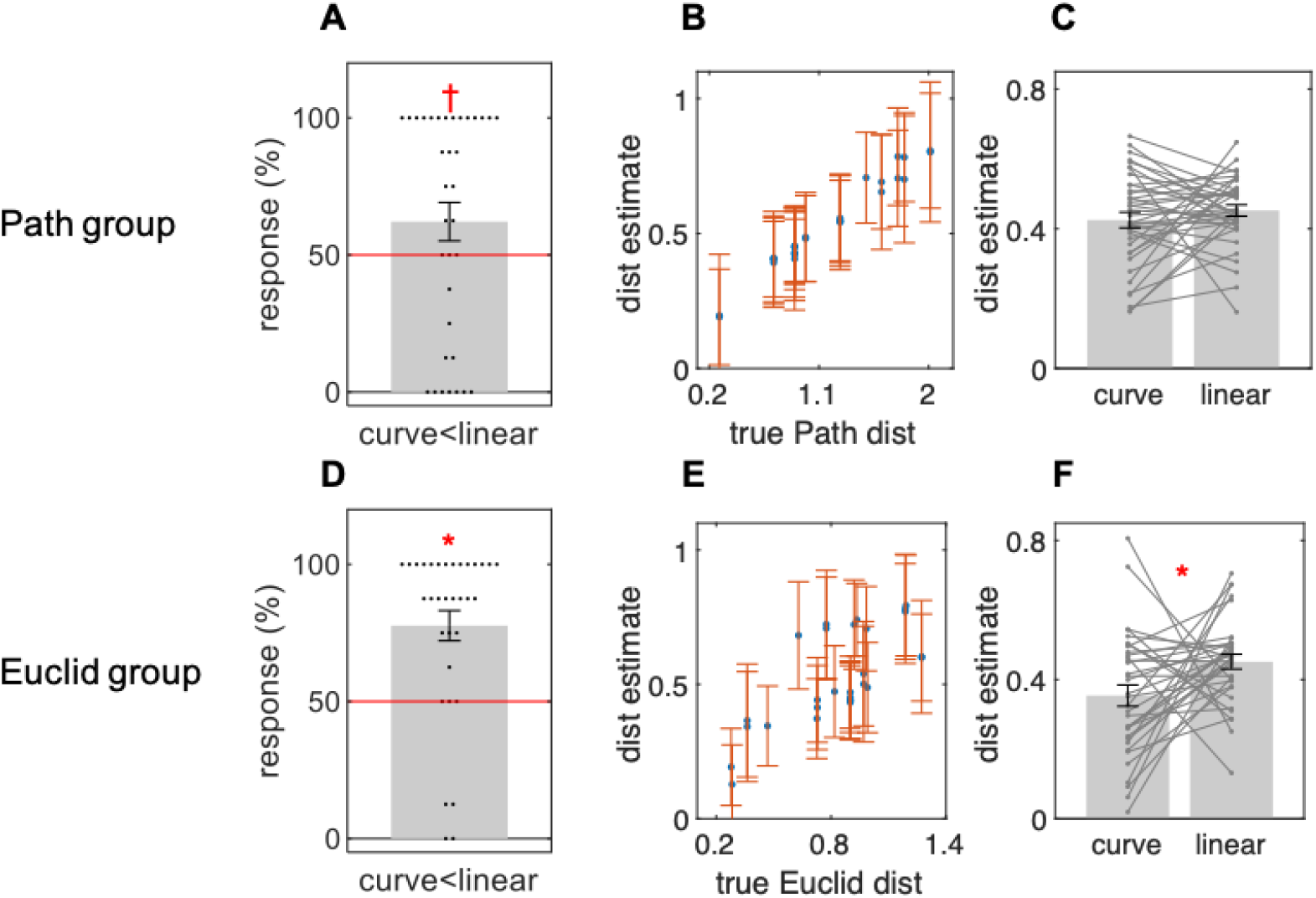
Distance estimate results in Experiment 2 (driving/looking). **A**. In the comparison task, participants showed a tendency for underestimating the path distance on the curve (p = 0.051). Black dots = individual participant; red line = chance level. **B**. Participant’s path distance estimates were highly correlated with the true path distance. Each dot = group mean distance estimate for unique object pair. **C.** The mean path estimate for the curve was not significantly different from the linear one. Dots = individual participant’s distance estimate between 0 (“very close”) and 1 (“very far”). **D**. Participants correctly reported the curve distance as shorter than linear during the comparison task. **E**. In the slider task the estimated distances were positively correlated with true Euclidean distance. **F**. Curved distance estimate was significantly shorter than the linear distance. All error bars are group SE.

For the debriefing question on the distance estimation strategy, 37% selected the option that “I imagined myself moving from one picture to another picture and estimating the distance (first-person perspective)” (Fig. 1C), 45% selected “I had a map of the 3D curved environment in my mind and measured the distance on that map (bird’s-eye view 3D)” (Fig. 1A), and 18% selected “I had the surface map of the environment, rather 2D, and measured the distance on that map (bird’s-eye view 2D).” (Fig. 1B).

#### 3.3.3 Distance estimation: Euclidean group

Most participants performed the distance estimation task well. Out of 35 individuals, 33 showed above chance accuracy for easy trials in the comparison task (mean accuracy for all participants = 92 ±14%). These participants correctly reported the curve as shorter in the comparison task (78 ±31%, sign-rank test, p < 0.001, Z = 3.5, 1-tailed, Fig. 4D). Participants’ distance estimates during the slider task were positively correlated with true Euclidean distance (ρ = 0.49 ±0.26, Fig. 4E). The curve distance estimate was also significantly shorter than the linear estimate during the slider task (mean estimate for the curve = 0.35 ±0.18 vs. linear = 0.45 ±0.12, t(34)=2.3, p = 0.014, one-tailed, Fig. 4F).

### 3.4. Discussion

In the current experiment, participants learned the object locations on the surface while driving as in the previous experiment, but additionally they could look above or below. We tested whether this additional viewing behavior encouraged participants to build a more precise 3D map of the environment, even though they were still physically constrained to the surface. If that were to be the case, we would expect more precise estimates of Euclidean distance, when they were explicitly asked (Euclidean group). It might also be the case that their path distance estimation would be strongly influenced by the 3D map, leading to an underestimation bias for the curve path (Path group).

When participants estimated the path distance, we found a tendency for underestimation of the curved path, similar to what we already observed in the previous experiment when participants were driving on the surface without looking up or down. This suggests that participants’ maps of the space were not completely flattened (schematic representation in Fig.1B) although such a flattened or topographic map could be the most efficient and adequate for the path estimation task. Similar to experiment 1, only a minority of participants reported that they imagined a flattened map during the distance estimation strategy. However, the influence of the 3D map was weak, only being observable in the direct comparison task, and not the slider task.

When participants estimated the Euclidean distance, the curve was reliably reported as shorter than the linear distance in both the comparison and slider tasks. Previously, we only found a significant difference between the curved and linear surfaces in the direct comparison task, and not in the slider task, which might be less sensitive to detect small differences between distances. This result fits our prediction that viewing behavior helps participants to build a global 3D representation of the environment. However, a direct comparison between the Euclidean distance estimate between the previous experiment and the current experiment didn’t yield a statistically significant difference (t(70)=1.1, p=0.3, Supplementary Fig. 7). Also, we note that the participants’ Euclidean distance estimation was still not perfect. First, the estimated ratio between the curve and linear distance in the slider task (median curve/linear = 0.79) was noticeably larger than the true ratio (0.48). Although, we should bear in mind that the distance estimate reported in the slider task does not necessarily have a strict linear relationship to the true distance. Second, when all pairs of between-object distances were considered, the correlation between the estimated and true Euclidean distance (ρ = 0.49 ±0.26) was significant, but not as strongly as the path estimation group who showed a very strong correlation (ρ = 0.65 ±0.34). However, it should be considered that Euclidean distance estimation can be inherently more difficult to report than the path distance because the range of the true Euclidean distance was smaller than the range of path distance (e.g. the maximum path distance between the objects was 2.0 whereas the maximum Euclidean distance was 1.3). Further, there was an additional confounding factor of the curved surface acting as a natural physical barrier, rendering the straight Euclidean distance estimation difficult (e.g. participants could not directly see location B when they were standing at location G because the curved surface was not transparent, Fig. 1A). As we already discussed in Experiment 1, barriers are known to deteriorate the distance estimation (Newcombe & Liben, 1982).

In sum, we again found that a completely flattened map was not utilized, although only the location and distance on the surface was behaviorally relevant. Participants could reliably distinguish the Euclidean distance on the curve and linear section, but the Euclidean distance estimation performance still seemed suboptimal. Might this mean that humans are inherently poor at building a 3D model of the world and distance estimation? In the following experiment we asked whether participants would be better at estimating Euclidean distance if they could freely fly in the virtual environment.

## 4. Experiment 3

### 4.1. Introduction

In Experiment 3, we removed the constraints of driving on the surface for participants. The objects that participants had to learn were still lying on the surface as in the previous two experiments, however, participants explored the environment with a flying motion. In contrast to Experiment 1 and 2, participants could directly move between the two points in 3D space along the straight line, as long as there was no natural barrier in the way. For example, a participant could fly along the nearly shortest (Euclidean) route from A to D. They still needed to take a small detour around the edge of the curved surface if they wanted to fly from A to B (Fig. 5). The flying route from A to B was still significantly shorter than the equivalent driving route on the surface. In this flying setup we wanted to test whether participants would build a more volumetric representation of the environment and show better Euclidean distance estimation. Therefore, we only tested Euclidean distance, not path distance on the surface, in this experiment.

**Figure 5.**
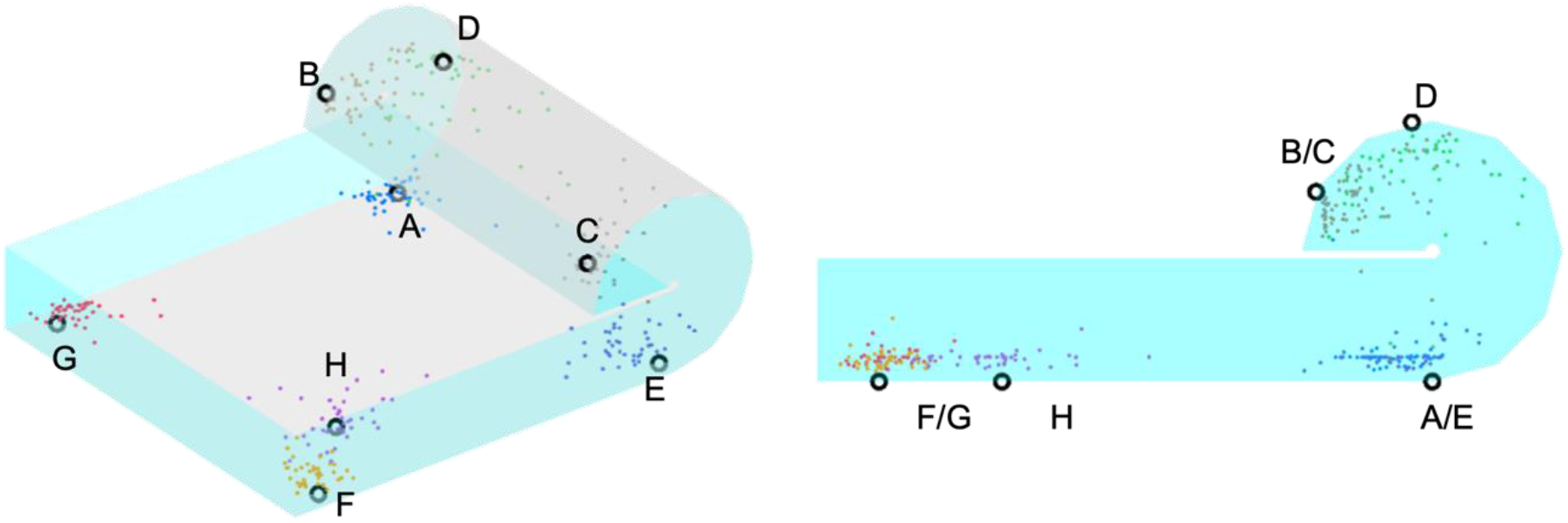
Object-location memory test result in Experiment 3. Remembered object locations in the last trial of all participants are shown in 3D view (left) and side view (right). Overall, remembered locations (color dots) were very close to the true location (black circles), but the errors were larger for the locations (B, C, D) on the curve side than the locations on the flat side (F, G, H).

### 4.2. Method

#### 4.2.1 Participants

Participants were recruited from the same online platform (www.prolific.com). Forty-one participants (female = 11, mean age = 23.5 ±4.5 years) completed the experiment. None of them took part in the previous experiments.

#### 4.2.2 Virtual environment

The identical environment was used as in the Experiment 1 and 2.

#### 4.2.3 Task and analysis

Participants completed the familiarization, object-location learning and test, two types of the distance estimation tasks, and the debriefing. The object locations, starting location and heading of the participants, trial structure, and experimental sequences were all identical to the previous experiments. The key change from the previous experiment was that participants could rotate, not only horizontally around the longitudinal body axis (yaw rotation), but also vertically around the side-to-side axis (pitch rotation). They could then move forwards or backwards in the direction the were facing. For instance, if they looked 45° upwards and pressed the forward movement key, they would move 45° upwards while maintaining the tilted heading. An example trajectory and a participant’s view in the flying condition can be found in <u>Supplementary Video 5</u> and Supplementary Fig. 6. Of note, there is one more degree of freedom for rotation in 3D, called the roll rotation, which allows agents to rotate around the front-to-back axis (e.g. left ear down, right ear up). However, controlling all 3 rotation axes using a keyboard or mouse with desktop-based VR is complicated and the roll rotation does not affect the movement direction. It is also known that humans tend to keep upright posture with zero roll (Barnett-Cowan & Bülthoff, 2013) and even flying bats have only a small number of cells tuned to roll rotation (Finkelstein et al., 2015). Thus, we only included the yaw and pitch components of rotation.

Controlling 3D rotation and flying movement can be more difficult compared to controlling driving motion in the previous experiments, so we allowed more time for the familiarization phase and object-location learning and test phase in the virtual environment (the time limit was set to 120 seconds from 60 seconds). We also increased the maximal number of repetitions per object from 8 to 10 during the object-location memory test. We do not think that this modification in a maximum number of repetitions per objects or time limit provided an advantage in object-location learning in the current experiment compared to previous experiments, because as we describe in the following result section (Section 4.3.1), the mean distance error and the mean number of repetitions was comparable to previous experiments. The time required for movement practice in the familiarization phase (201 ±70 seconds) and the mean trial duration during the object-location memory test (12.1 ±5.9 seconds) was longer than the previous driving experiments (c.f. mean familiarization duration = 126 ±57 seconds, mean object-location trial duration = 8.5 ±3.1 seconds, Experiment 1 and 2 combined), probably due to difficulty of controlling flying motion.

### 4.3. Result

#### 4.3.1 Object location memory

Similar to the previous driving experiments, most participants placed all objects in the close vicinity of the correct location (mean repetition per object = 5.8 ±1.3, mean distance error = 0.10 ±0.05 for all 41 participants, all distances are reported as relative to the short axis of the environment, Fig. 5). Further, only those who remembered each of the 8 objects within a specific learning criterion (Euclidean distance error of max. 0.25) were included in the analysis, leaving n = 35. The final displacement error was larger for objects at the curved part compared to objects on the flat part (error for the objects on the curve = 0.10 ±0.03 vs. flat = 0.07 ±0.02, t(34) = 5.2, p < 0.001). This was not surprising because larger displacement error was possible along the curved surface, when participants were flying, due to their oblique trajectories toward the targets, whereas the objects on the flat surface could be still approached as if participants were driving with near zero height from the plane (Supplementary Fig. 6).

#### 4.3.2 Euclidean distance estimation

Similar to the previous experiments, most participants passed the manipulation check (easy trials) for the distance estimation task (34 out of 35). The accuracy for comparing the curve and linear trials was also high (82 ±28%, sign-rank test, Z = 4.4, p < 0.001, Fig. 6A). In the slider task, participants showed a good fit between the estimated and correct Euclidean distance (ρ = 0.49 ±0.26, Fig. 6B). The curve distance was estimated as significantly shorter than the linear distance (curve estimate = 0.35 ±0.15 vs. linear = 0.57 ±0.16, t(34) = 5.2, p < 0.001, Fig. 6C). It is noteworthy that the magnitude of the difference between the curve and linear distance estimates in this experiment was significantly larger than the difference observed in the previous experiments with driving (1-way ANOVA, F(2,104)=5.3, p=0.007, post-hoc pairwise comparison, Exp1 vs. Exp3, t(70)=3.3, p=0.002 (uncorrected); Exp2 vs. Exp3, t(68)=2.0, p=0.04 (uncorrected), Supplementary Fig. 7). Further, the curve/linear ratio was comparable to the true Euclidean distance ratio (true = 0.40, the current flying experiment = 0.57, previous driving/looking experiment = 0.79).

**Figure 6.**
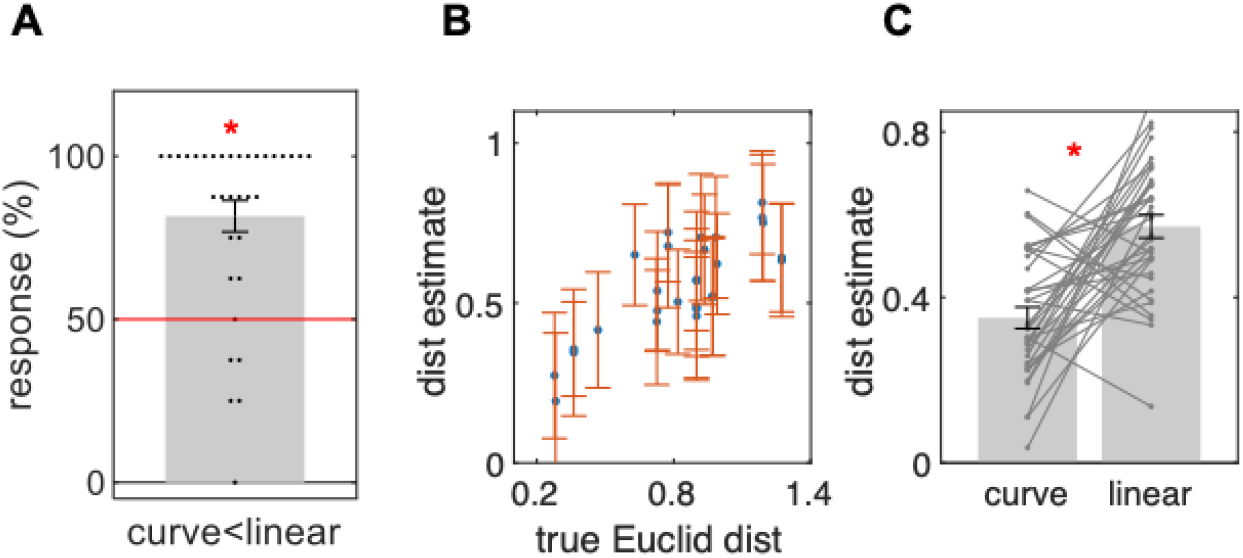
Distance estimation result in Experiment 3 (flying). **A**. Participants showed high accuracy for the comparison task. **B**. In the slider task, the estimated distances were positively correlated with true Euclidean distance. **C**. Curved distance estimates were significantly shorter than the linear distance. All error bars are group SE.

### 4.4. Discussion

As we hypothesized, participants who explored the environment with flying motion were better at estimating the Euclidean distance, compared to the previous experiments where participants explored the environment with driving motion. We believe that the experience of 3D rotation and movement encouraged participants to build a more appropriate 3D model of the environment, enabling a better distance estimation. Distance estimation could also have directly benefited from temporal memory of the travel distance. Humans can estimate distance using both static information (e.g. visual, auditory depth cues) and dynamic information (e.g. optic flow, self-motion)(Sun et al., 2004). In the previous driving experiments, participants could estimate the Euclidean distance on the curved surface using only the static depth cue or abstract representation of the environment, whereas the flying group could also recall the temporal duration (of getting from point A to point B) while estimating the Euclidean spatial distance. It is known that some people rely more on an implicit, time-based method during spatial distance estimation tasks (Mossio et al., 2008).

Of note, we noticed that the overall correlation between the estimated and true Euclidean distance, for all pairs, was significantly positive but still not as strong as between the estimated and true path distances in the previous driving experiments. This implies that there is still room for improvement in the Euclidean distance estimation. We think that the transparent wall surrounding the surface and the curved surface itself rendered the Euclidean distance estimation more difficult for particular pairs of objects such as G and B or G and D. To move between these pairs, participants had to take a detour around the wall (Supplementary Fig. 6). It has been shown that physical or contextual boundaries can divide space into compartments, hindering Euclidean distance judgments (Han & Becker, 2014; Hirtle & Jonides, 1985).

## 5. General discussion

Humans and most mammals dwell and navigate on 2D surfaces embedded within the 3D world. Terrains can have rather complex 3D profiles with bumps and holes, but a vast majority of previous research on spatial cognition was focused on horizontal flat surfaces. The 2D Cartesian map has become the predominant cognitive map in neuroscience research. In the current study we used a novel 3D virtual environment containing a curved surface to test whether humans represent such an environment using a 2D flattened map or a more volumetric 3D map. While a few previous studies have investigated participants’ sense of orientation in 3D space using a maze consisting of narrow vertical and horizontal tracks (Indovina et al., 2016; Vidal et al., 2004, 2006), slanted surfaces (Nardi et al., 2011; Restat et al., 2004; Steck et al., 2003), or a multilevel building (Brandt et al., 2015; Kim & Maguire, 2018; Montello & Pick, 1993), spatial representations of large, navigable, curved surfaces remained scarce.

The first main finding of the current study was that participants were aware of the global 3D layout of the environment despite their movement being restricted to the surface and despite the fact that the object-location memory task could be solved with 2D coordinates on the surface (Experiment 1 and 2). This implies that humans do not necessarily extract the pure topographic, relational knowledge (e.g. links between the key object locations) and store them in a 2D Cartesian map. Rather, they also hold information about the 3D world. It might seem somewhat suboptimal from a computational perspective, considering that the brain can extract a low-dimensional structure in a complex high-dimensional world, e.g. a principal component of high-dimensional stimulus space (Chang & Tsao, 2017; Summerfield et al., 2020). However, the automatic encoding of a 3D layout could be the natural behavior given the saliency of 3D cues and the potential benefit of acquiring 3D knowledge for future navigational problems. This might be related to the ability of animals to take 2D vector shortcuts after only following 1D routes (Tolman, 1948). Previous behavioral experiments have shown that people use the vertical axis of a slant surface as an orientation cue to facilitate their spatial memory (Nardi et al., 2011, 2021; Restat et al., 2004; Steck et al., 2003). Importantly, participants in our studies did not have any vestibular inputs, nor did they experience energy cost along the curved surface because we let participants move with constant speed in our virtual environment as if there is no gravity, in contrast to abovementioned experiments where participants experienced the energy cost when moving upward along the slope in the real world. The absence of vestibular inputs and an artificial environment is a limitation of the current study, but we could learn that the visual cues alone (e.g. sky, mountain landscape) are sufficient to trigger automatic encoding of the vertical or orthogonal axis to the surface of locomotion.

If the participants did not build a completely flattened 2D map, did they instead build a metric 3D map, containing precise Euclidean distance and angle information between the locations? This is also unlikely given the suboptimal 3D Euclidean distance estimation performance, even in the flying condition (Experiment 3). We propose a mixture of a metric map and topographic knowledge as the most likely form of spatial representation. Originally, O’Keefe and Nadel made a strong claim that the cognitive map is Euclidean (O’Keefe & Nadel, 1978). However, many studies have since challenged the notion of a strict metric or Euclidean map. Behaviorally, Warren and colleagues have shown that participants successfully navigate in a virtual environment where invisible wormholes break the rules of 2D Euclidean geometry (Ericson & Warren, 2020; Warren et al., 2017). Interestingly, the participants did not even notice the peculiarity of the environment, ruling out the necessity for Euclidean map knowledge. Neurally, hippocampal place cells change their receptive fields when the environment changes its shape and size by stretching and shearing, but preserve the relative location or topological information about the environment (Dabaghian et al., 2014; O’Keefe & Burgess, 1996). Again, these results imply that space is not encoded in the brain like a precise Cartesian coordinate system with exact distance and angle from an origin. Alternatively, the model of the environment can be graph-like, consisting of nodes and links, and metric information about distance and angle between the nodes or landmarks is locally embedded, rather than forming a globally coherent 3D metric map (Peer et al., 2020; Warren, 2019). Combining the graph knowledge and local metric information can be used in robotics and artificial agents (Hübner & Mallot, 2007; Mallot & Basten, 2009). On a related note, Glennerster proposed that an observer in a 3D world does not need to reconstruct a 3D scene, rather, they could use the representation somewhere in between the 3D reconstruction and a more 2D image-based representation, which is updated upon movement of the observer in space (Glennerster, 2016). For instance, translation and rotation of vantage points provides information on the distance, slant, and depth of surfaces without necessarily reconstructing the perfect 3D scene. In our experiment, the most likely encoding scenario would be to remember the fine-scale relative location on the surface while holding rather coarse information about the layout of the surface within a global 3D world.

When we study the mental model of the world used by people, we should not underestimate the importance of how they explore the environment. In the present study, participants explored the identical 3D environment with varying modes of exploration, from driving alone (Experiment 1) to driving with vertical viewing (Experiment 2), to flying (Experiment 3) and we observed the enhanced 3D Euclidean distance estimation performance with the increased degree of freedom in 3D movement. From an ecological psychology perspective, active interaction between the observer and the environment should be given more importance than a static and rigid representation of the external world (Costall, 1984). Previous research in 2D flat environments has shown that the accuracy of spatial memory and shortcut behavior is dependent on which perspective participants took while learning the environment (1st-person-perspective, birds eye view, or hybrid slanted perspective) (Barra et al., 2012). At the neural level, the firing pattern of place cells is not only determined by the location and physical environment but also by trajectories and goal locations (Grieves et al., 2016). Therefore, it is crucial to consider the complexity of exploratory behavior and its ecological validity when we want to understand the mental representation of the external environment, be it flat or curved, 2D or 3D.

In conclusion, our study provided novel insights on human spatial memory and mental map formation of curved surfaces embedded within a 3D world. The cognitive map is neither completely reduced to 2D nor is it fully 3D. Rather, it is somewhere in between. We believe that the representation of the environment is flexible and adapted to behavioral experience and demand, such as how participants are interacting with the environment (e.g. driving or flying) and which type of spatial information needs to be recalled (e.g. object location, path, or Euclidean distance). Furthermore, our study encourages investigation of more general cognitive maps beyond 2D Euclidean space. It has been proposed that the neural mechanisms encoding navigable space, such as place codes and grid codes, can serve as general coding principles for encoding abstract knowledge space (Behrens et al., 2018; Bellmund et al., 2018; Theves et al., 2019, 2020). Despite non-physical space having many more dimensions, not all of which are independent from one another, previous literature has focused 2 dimensions. These ‘abstract spaces’ have consisted of 2 orthogonal feature dimensions such as two independent smells, lengths of visual features, or personal characteristics (Bao et al., 2019; Constantinescu et al., 2016; Park et al., 2020; Tavares et al., 2015; Viganò & Piazza, 2020), or alternatively, the conceptionally relevant 2D representation within a 3D feature space (Theves et al., 2020), which is analogous to a 2D flat surface. In due course, we hope to gain a better understanding of how humans develop their internal representation of high-dimensional, non-physical space, which contains a low-dimensional structure, such as the cognitive map for a 2D curved surface within a 3D abstract world.

## Data and code availability

All data and analysis scripts are available in OSF (https://osf.io/gsnyx/).

## Author contributions

MK and CFD developed the study concept and wrote the manuscript. MK programmed the experiment, collected, analyzed the data.

## Acknowledgement

This work has been supported by the Max Planck Society. CFD’s research is further supported by the European Research Council (ERC-CoG GEOCOG 724836), the Kavli Foundation, and the Jebsen Foundation.

## Declaration of interest

Authors declare no competing interests.

## Supplementary Materials

Supplementary Videos can be found in OSF repository, which also contains raw data and analysis scripts. Link and caption for each Supplementary Videos are attached below.

**Supplementary Video 1.** Familiarization period during Experiment 1. Participants drove on the surface with their head parallel to the ground. They followed the traffic cone and practiced the movement. Guide arrows were shown on the ground to help them quickly find the target.

https://osf.io/7qfvg/

**Supplementary Video 2.** Instruction video for Euclidean distance estimation task.

https://osf.io/bp4q6/

**Supplementary Video 3.** Video for the convex surface experiment.

https://osf.io/zfsbt/

**Supplementary Video 4.** Familiarization period during Experiment 2. Participants drove on the surface and they could also rotate their views up and down.

https://osf.io/4tep3/

**Supplementary Video 5.** Familiarization period during Experiment 3. Participants were not restricted to the surface and they could rotate on yaw and pitch plane and move forward or backward as if flying in the air.

https://osf.io/82hqf/

**Supplementary Figure 1.**
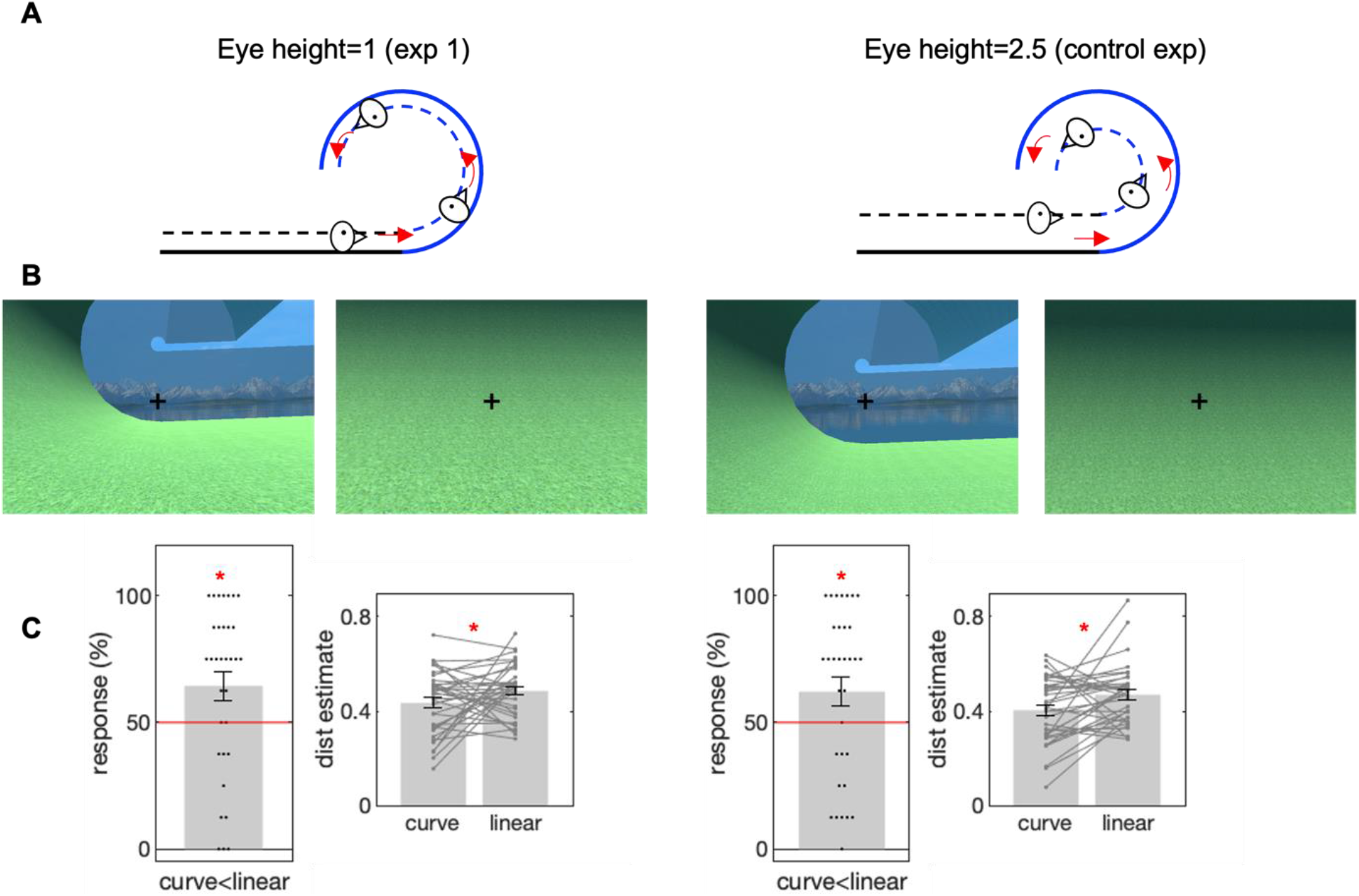
Manipulation of eye height in the control experiment. **A**. Physical path length on the curved (blue solid line) and the linear (black solid line) part of the surface were identical, but the path length at the eye level was shorter on the curved section (blue dashed line) compared to the linear section (black dashed line), and this difference was greater when the eye height was higher. **B**. Example views from the same locations with different eye heights. Black cross at the center of the screen was shown here to help readers to see the difference between the eye heights (e.g. eye height was about 20% of the radius in the Experiment 1 and about 50% of the radius in the control experiment). The cross was not shown during the actual experiment. **C**. The magnitude of the underestimation biases for the curved path was similar across the two experiments of different eye heights. In the comparison task, the mean rate of curved section chosen as shorter was significantly above chance in both experiments (Experiment 1, 64 ±33%, n = 33, Z = 2.2, p = 0.013; control experiment, 62 ±33%, n = 33, Z = 2.0, p = 0.023, sign-rank test, one-tailed) and the difference between the two experiments was insignificant (ranksum test, p=0.8). The slider task also showed the similar underestimation bias in both experiments (the difference between the curve and path, Experiment1, −0.050 ±0.18, t(36) = 1.7, p = 0.045, one-tailed; control experiment, −0.066 ±0.17, t(35)=-2.4, p=0.011, one-tailed; the difference between the two experiment, t(71)=0.38, p=0.72). Of note, the result for the Experiment 1 was already reported in the main text (Fig. 3C,E) but shown here again for the ease of comparison between the two experiments. All error bars are group SE.

**Supplementary Figure 2.**
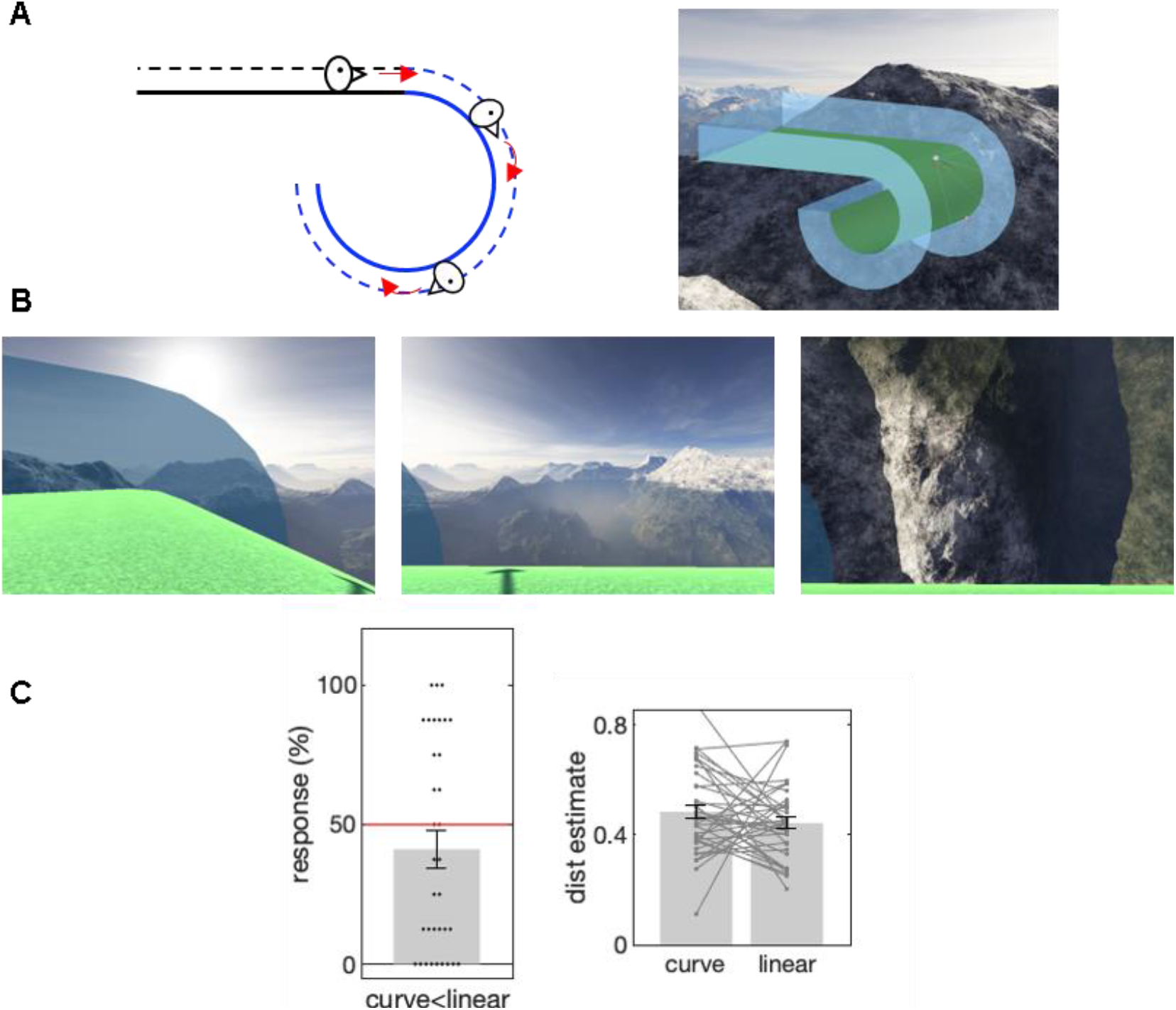
A control experiment where participants drove outside of the curved surface (“convex”). **A**. In contrast to the main experiments where participants drove inside of the curved surface (“concave”), the path length at the eye level was longer on the curved section (blue dashed line) compared to linear section (black dashed line). Please note that, driving on the convex surface induced strong sense of falling down and potentially anxiety, compared to the main experiment. Also, 3D Euclidean shortcut can be more difficult to imagine because such shortcut has to pass through the solid ground, unlike in the concave surface. **B**. Example views on the convex condition. Participants mostly see the upside-down mountainous landscape when they drove along the curved surface (middle, right panel). This was in a stark contrast to the main experiments where participants mostly saw the ground surface. **C**. There was no significant underestimation biases for the curved path. If anything, we observed a trend for overestimation bias in the comparison test, the curve path chosen as the shorter one was 41±38% (n = 34, Z = −1.5, p = 0.13). The difference between the curve and linear path estimate in the slider task was also not significant (curve: 0.48 ±0.16, linear: 0.44 ±0.13, t(36)=1.2, p=0.26). All error bars are group SE.

**Supplementary Figure 3.**
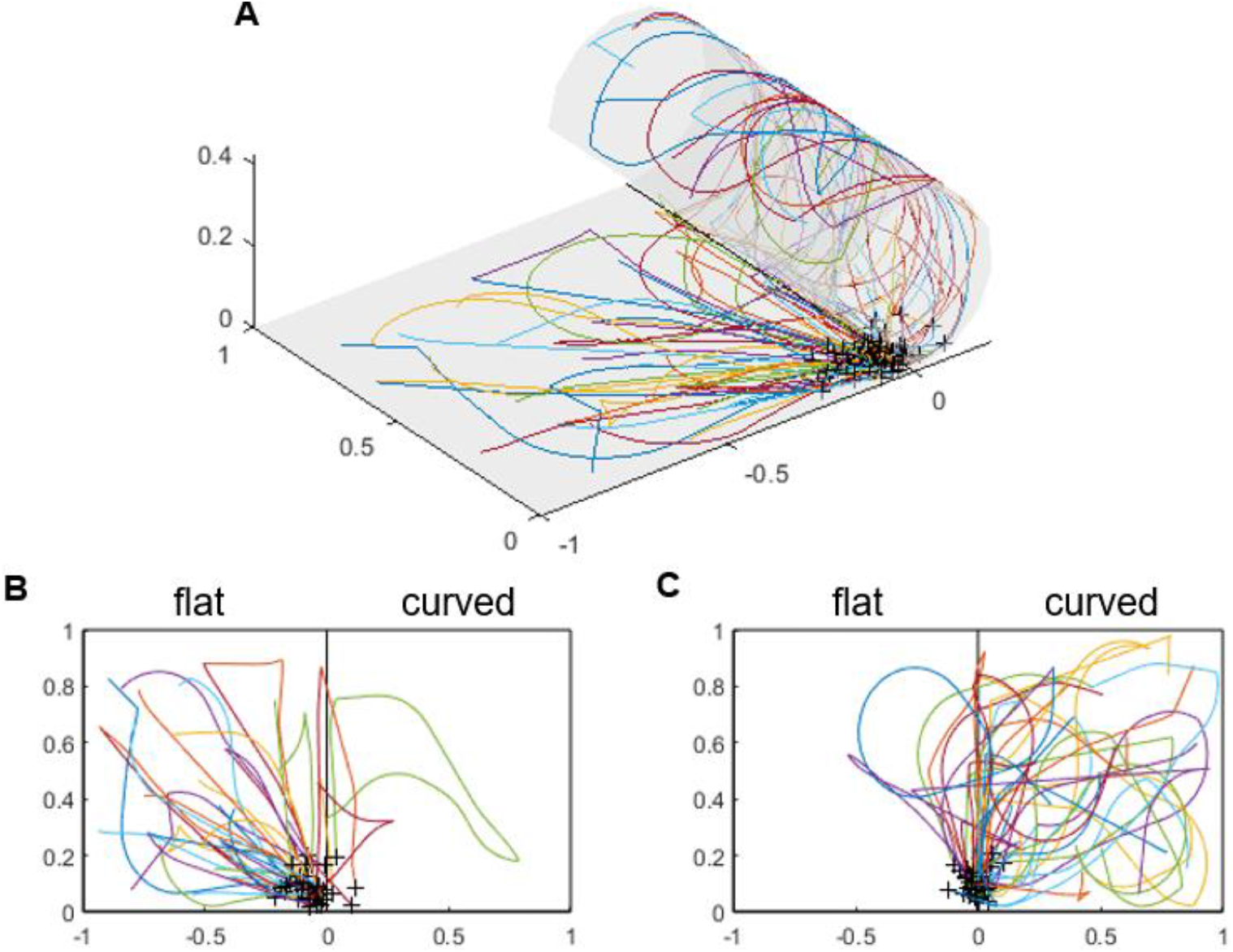
Trajectories during the object-location test in Experiment 1. **A**. The trajectories of all participants (n=83) who correctly placed the middle object E (true location: [0, 0.05] in the normalized 2D coordinate) in the last trial are overlaid on 3D surface. Each colored line indicates participants. Black cross indicates the drop locations (remembered location). The distribution of drop locations was skewed towards the flat part of the environment. We separately show the trajectories in which participants started from the flat (n=41) and curved (n=42) part of the environment and overlaid them on the flattened surface in the panel B and C, respectively. **B**. Participants tended to move straight from the start location on the flat side to the correct target location at the midline, and they mostly did not pass the midline. **C**. Participants who started from the curved part were more likely to pass the midline or take a small detour and approach the midline object from the flat part.

**Supplementary Figure 4.**
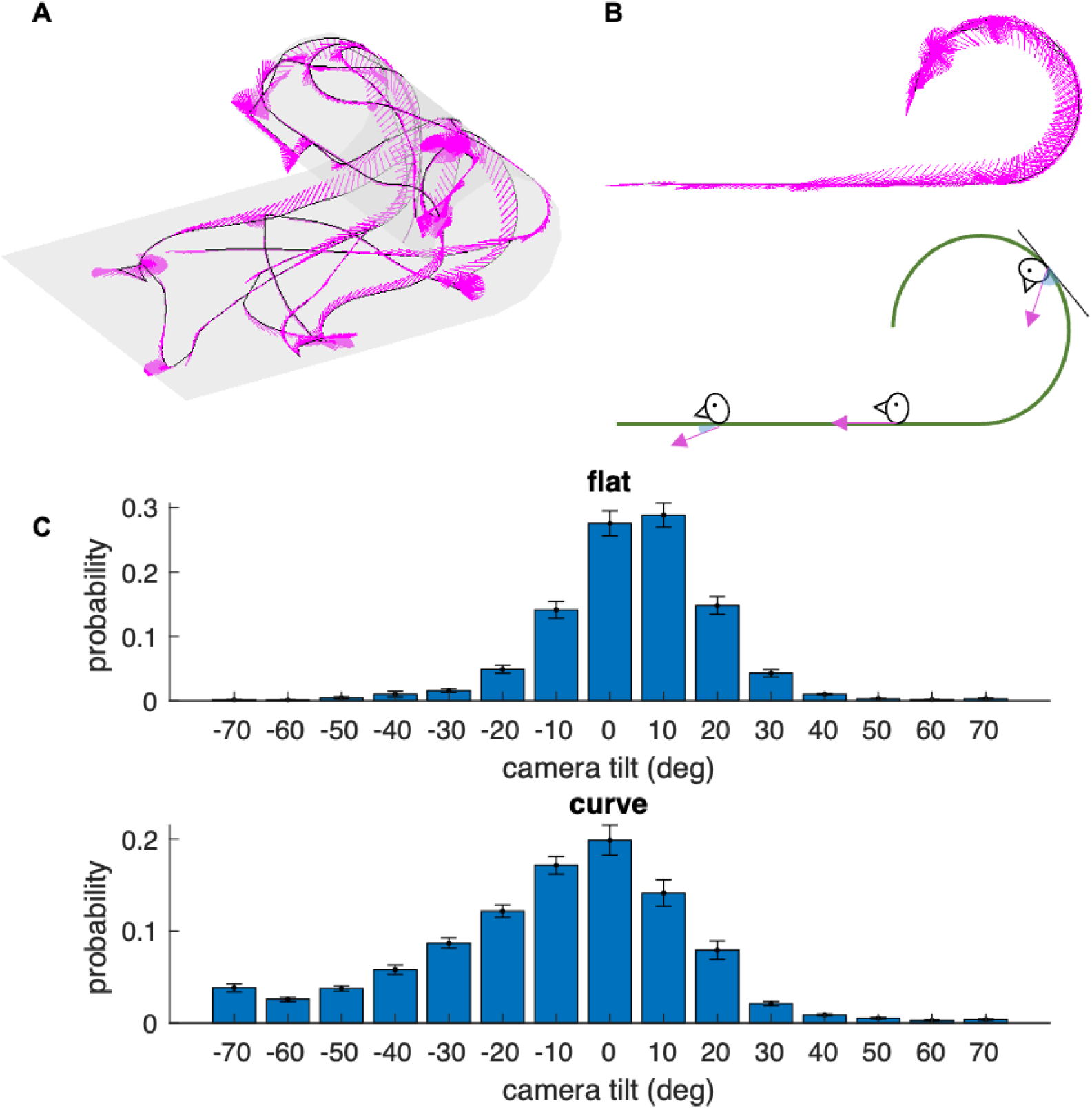
Viewing behaviors in the Experiment 2. **A**. One example participant’s trajectory during the familiarization period is shown. The black line shows the location of this participant. Facing directions of the camera were visualized at regular time intervals with magenta lines. When a participant moved straight with a zero vertical tilt of camera, the camera facing direction (magenta) and the trajectory (black) overlap. When a participant stood still and rotated, camera facing directions (magenta) form pie shapes. **B**. The vertical tilt of the camera can be readily shown from the side view. Top, camera facing directions of the same participant are shown as magenta lines; bottom, schematic view. On the flat part of the environment, participants mainly looked straight ahead and the camera direction (magenta arrow) was parallel to the ground (tilt = 0) or they looked slightly towards the surface (tilt > 0); in contrast, participants often looked away from the surface (tilt < 0) on the curved part, therefore the camera facing direction (magenta arrow) angled away from the tangent of the surface (black line). **C**. The distribution of camera tilts on the flat and curved part of the environment for all participants during the object-location test phase. Error bar, group SE.

**Supplementary Figure 5.**
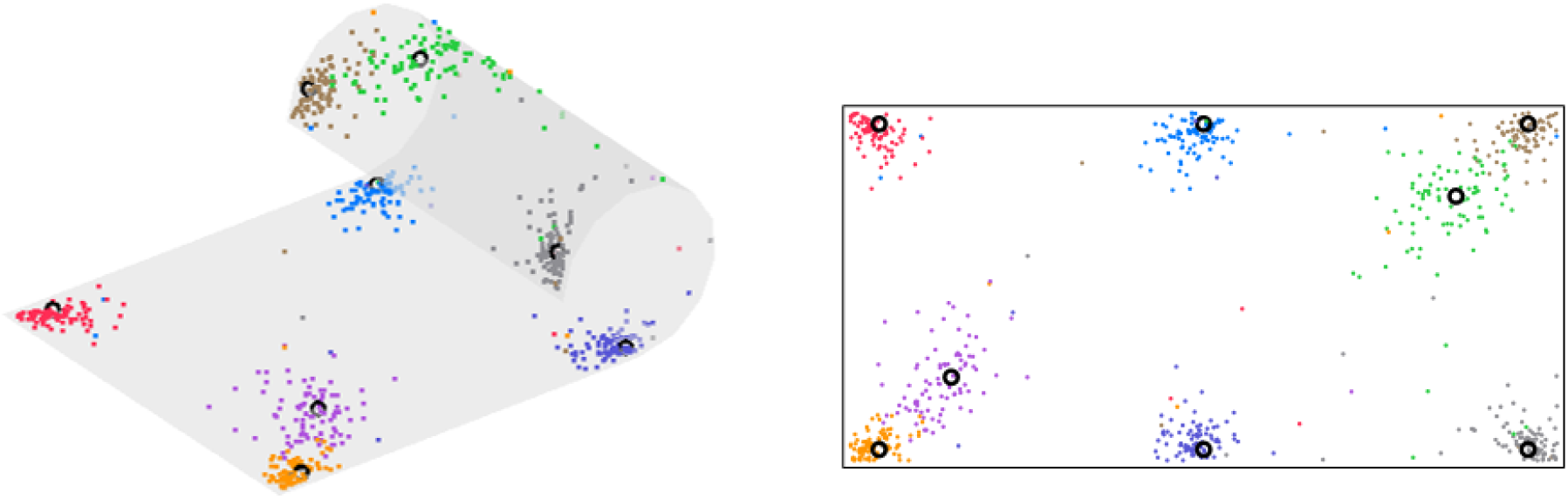
The object-location memory test result in Experiment 2. Similar to Experiment 1, most participants remembered the location well at the end of the test phase (mean distance error < 0.1). Color dots, the last drop locations of objects for all participants; black circle, the true location. Left, 3D view; right, flattened view. Distance was normalized to the length of the short axis of the surface.

**Supplementary Figure 6.**
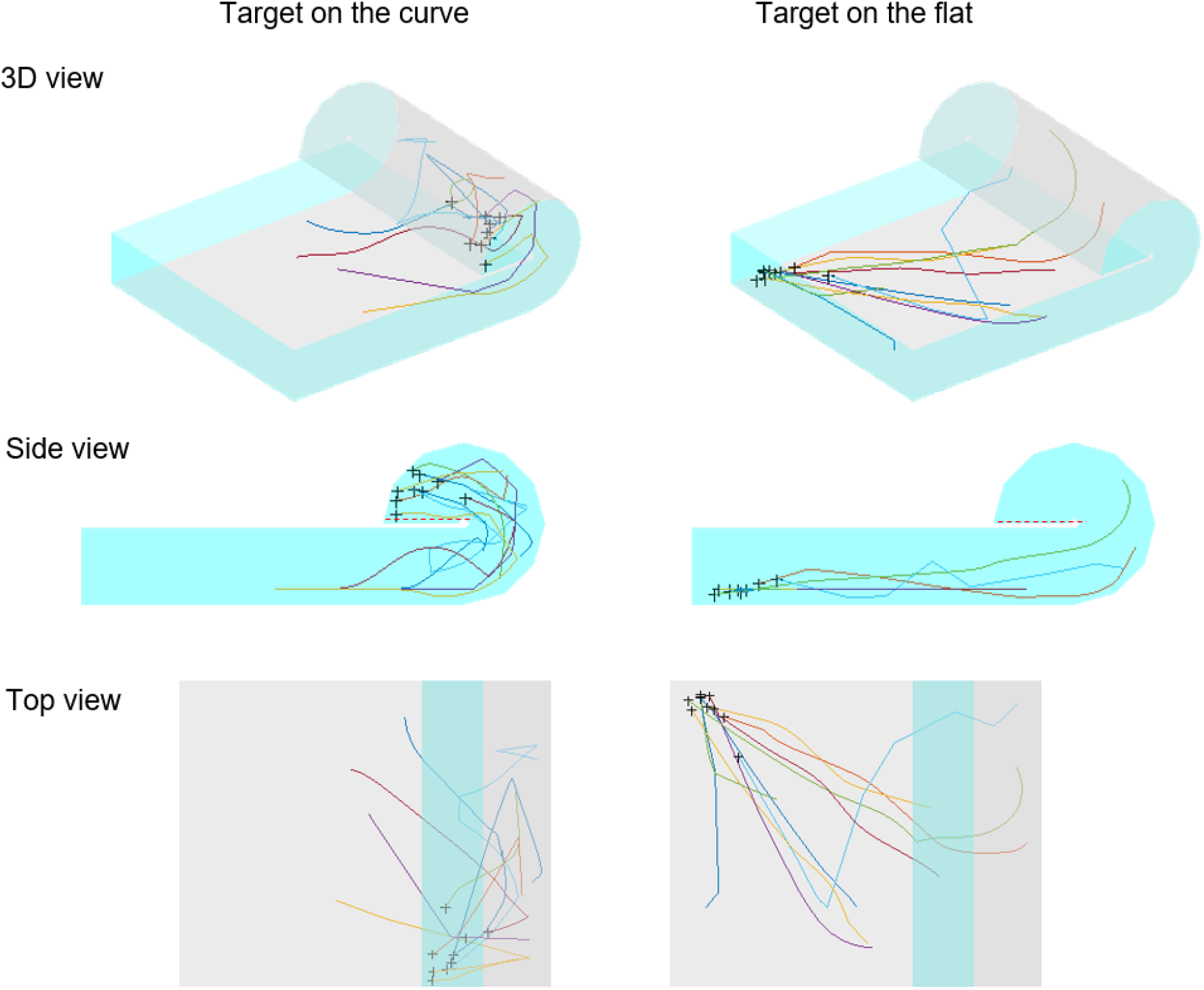
Movement trajectories during the object location test in Experiment 3. Ten randomly selected participant’s trajectories toward the target location on the curve (left column) and on the flat (right column) part of the environment are shown. When the target location was on the curved surface, participants had to take a turn around the transparent wall at the end of the curve (red dash line on the side view). Participants could move almost parallel to the ground when they approached the location on the flat part. Each color line represents each participant and the black cross shows the end location of the trajectory.

**Supplementary Figure 7.**
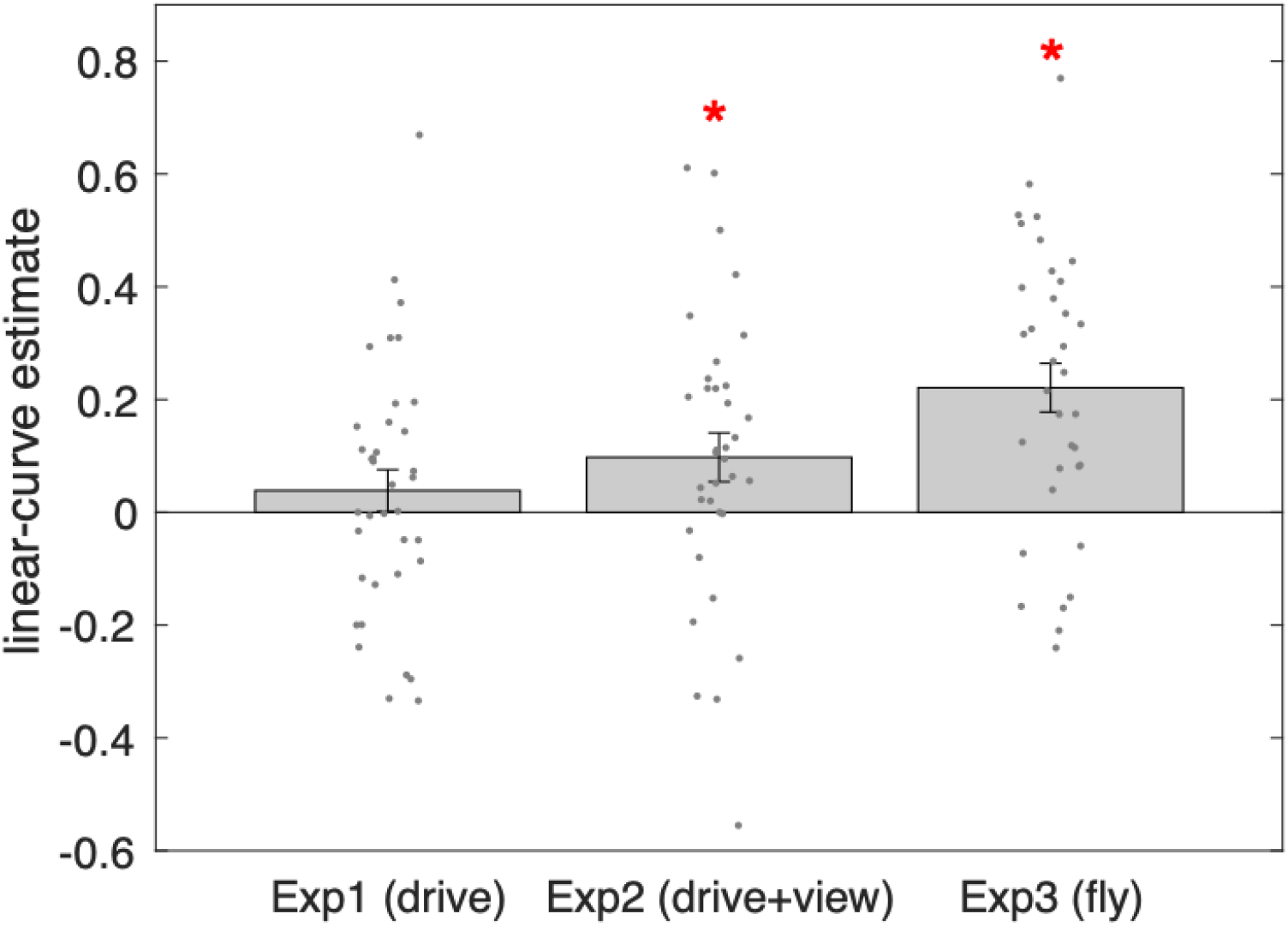
A summary of the Euclidean distance estimates during the slider task for three experiments. In Experiment 2 and Experiment 3, participants reliably distinguished the linear and curve section (one-sided t-test against zero, Exp1, t(36)=1.1, p=0.15; Exp2, t(34)=2.3, p=0.014; Exp3, t(34)=5.2, p<0.001). The distance estimate difference was greatest in the Experiment 3 where participants could fly (1-ANOVA, F(2,104)=5.3, p=0.007, post-hoc pairwise comparison, Exp1 vs. Exp2, t(70)=-1.1, p=0.3; Exp1 vs. Exp3, t(70)=-3.3, p=0.002; Exp2 vs. Exp3, t(68)=-2.0, p=0.044). All error bars are group SE.

## Notes

https://osf.io/gsnyx/

### Competing Interest Statement

The authors have declared no competing interest.

### Summary of Updates

Control experiments with eye height manipulation and convex surfaces were added.

